# Identifying TCDD-resistance genes via murine and rat comparative genomics and transcriptomics

**DOI:** 10.1101/602698

**Authors:** Stephenie D. Prokopec, Aileen Lu, Sandy Che-Eun S. Lee, Cindy Q. Yao, Ren X. Sun, John D. Watson, Richard de Borja, Ada Wong, Michelle Sam, Philip Zuzarte, John D. McPherson, Allan B. Okey, Raimo Pohjanvirta, Paul C. Boutros

## Abstract

The aryl hydrocarbon receptor (AHR) mediates many of the toxic effects of 2,3,7,8-tetrachlorodibenzo-*p*-dioxin (TCDD). However, the AHR alone is insufficient to explain the widely different outcomes among organisms. Attempts to identify unknown factor(s) have been confounded by genetic variability of model organisms. Here, we evaluated three transgenic mouse lines, each expressing a different rat AHR isoform (rWT, DEL, and INS), as well as C57BL/6 and DBA/2 mice. We supplement these with whole-genome sequencing and transcriptomic analyses of the corresponding rat models: Long-Evans (L-E) and Han/Wistar (H/W) rats. These integrated multi-species genomic and transcriptomic data were used to identify genes associated with TCDD-response phenotypes.

We identified several genes that show consistent transcriptional changes in both transgenic mice and rats. Hepatic *Pxdc1* was significantly repressed by TCDD in C57BL/6, rWT mice, and in L-E rat. Three genes demonstrated different AHRE-1 (full) motif occurrences within their promoter regions: *Cxxc5* had fewer occurrences in H/W, as compared with L-E; *Sugp1* and *Hgfac* (in either L-E or H/W respectively). These genes also showed different patterns of mRNA abundance across strains.

The AHR isoform explains much of the transcriptional variability: up to 50% of genes with altered mRNA abundance following TCDD exposure are associated with a single AHR isoform (30% and 10% unique to DEL and rWT respectively following 500 μg/kg TCDD). Genomic and transcriptomic evidence allowed identification of genes potentially involved in phenotypic outcomes: *Pxdc1* had differential mRNA abundance by phenotype; *Cxxc5* had altered AHR binding sites and differential mRNA abundance.

**Author Summary:** Environmental contaminants such as dioxins cause many toxic responses, anything from chloracne (common in humans) to death. These toxic responses are mostly regulated by the *Ahr*, a ligand-activated transcription factor with roles in drug metabolism and immune responses, however other contributing factors remain unclear. Studies are complicated by the underlying genetic heterogeneity of model organisms. Our team evaluated a number of mouse and rat models, including two strains of mouse, two strains of rat and three transgenic mouse lines which differ only at the *Ahr* locus, that present widely different sensitivities to the most potent dioxin: 2,3,7,8 tetrachlorodibenzo-*p*-dioxin (TCDD). We identified a number of changes to gene expression that were associated with different toxic responses. We then contrasted these findings with results from whole-genome sequencing of the H/W and L-E rats and found some key genes, such as *Cxxc5* and *Mafb*, which might contribute to TCDD toxicity. These transcriptomic and genomic datasets will provide a valuable resource for future studies into the mechanisms of dioxin toxicities.

## Introduction

TCDD (2,3,7,8-<u>t</u>etra<u>c</u>hloro<u>d</u>ibenzo-*p*-<u>d</u>ioxin) is a member of the dioxin class of environmental pollutants. It is a persistent, highly lipophilic compound that can be created as a by-product during production of some herbicides and through the incineration of chlorine-containing compounds [1]. TCDD toxicity impacts almost all organ systems in mammals, with effects ranging from chloracne (particularly in humans) to immunosuppression, wasting syndrome, hepatotoxicity and acute lethality [2]. There are large differences in the lethality of TCDD, both between and within a given species. Two rodent species, hamster and guinea pig, show roughly 5000-fold difference in sensitivity to TCDD toxicity, with LD_50_ values of 1157-5051 μg/kg and 0.6-2 μg/kg, respectively [2]. The DBA/2 mouse strain exhibits a 10-fold lower TCDD responsiveness compared with the C57BL/6 strain to a wide variety of biochemical and toxic impacts of TCDD [reviewed in 3]. For example, the LD_50_ for male DBA/2 and C57BL/6 mice are 2570 and 180-305 μg/kg respectively [4,5]. Perhaps the most dramatic example of intraspecies differences in TCDD-susceptibility is the TCDD-resistant Han/Wistar (H/W) rat, which has an LD_50_ of >9600 μg/kg TCDD. By contrast, other rat strains are dramatically more TCDD-sensitive; the Long-Evans (L-E) rat has LD_50_ values of 9.8 μg/kg in females and 17.7 μg/kg in males [6].

Structural features of the aryl hydrocarbon receptor (AHR) play a major role in the diversity of TCDD-induced toxicities across species and strains. The AHR is a ligand-dependent transcription factor in the PER-ARNT-SIM (PAS) superfamily, which is evolutionarily conserved across fish, birds and mammals [7]. Normally bound to chaperone proteins including hsp90 and XAP2 in the cytosol, AHR can be activated by binding of ligands to the PAS-B domain, leading to nuclear localization [8]. Once in the nucleus and free of chaperone proteins, AHR dimerizes with the AHR Nuclear Translocator (ARNT) protein and binds to AHR response elements (AHREs) in the genome, altering transcription of specific target genes [9]. The AHR also has other, ‘non-genomic’ actions [reviewed in 10]. For example, ligand activation of the AHR leads to increased intracellular Ca^2+^, kinase activation and induction of Cox-2 transcription to promote a rapid inflammatory response [11].

Evidence for involvement of the AHR in TCDD toxicity comes from numerous studies, including species with structural AHR variants and *Ahr* knockout models. Mice lacking the *Ahr* gene are phenotypically [12,13,14] and biochemically [15,16] unresponsive to TCDD, as are *Ahr*-knockout rats [17]. Additionally, differences in the structure of the AHR protein result in a wide range of susceptibilities to dioxin toxicity. For example, the H/W rat is TCDD-resistant primarily due to a point mutation in the transactivation domain of the *Ahr* gene. This creates a cryptic splice site, leading to two distinct protein products (termed the deletion (DEL) and insertion (INS) isoforms) which are shorter than the wild type rat AHR (present in the L-E strain) [18]. Of these, expression of the INS isoform is predominant; however both are expressed in a number of tissues [19]. Numerous studies have sought to exploit these genetic differences among strains and species to decipher the mechanisms of dioxin toxicity [20,21,22,23].

The AHR is not the only mediator of dioxin toxicities: evaluation of rat lines generated through breeding of H/W and L-E rats suggests involvement of a second gene (termed gene “B”) in the extreme resistance of H/W rats to TCDD-induced toxicities [24,25]. Line A (Ln-A) rats contain the H/W *Ahr* and demonstrate similar resistance to TCDD-induced lethality as H/W rats; however, they appear to harbour the wild type (L-E) form of an as-yet unidentified gene “B” that is proposed to contribute to the phenotypic response to TCDD. Alternatively, Line B (Ln-B) rats express the wild type *Ahr* along with the H/W form of gene “B”, and demonstrate an intermediate LD_50_ of 830 μg/kg TCDD. Finally, Line C (Ln-C) rats do not fall far from L-E rats in sensitivity to TCDD (LD_50_ 20-40 μg/kg), and express wild type forms of both the *Ahr* and gene “B” [24]. It is thus unclear which responses are due solely to the various AHR and gene “B” isoforms and which are artifacts of the genetic heterogeneity at non-AHR loci among species and strains.

To isolate the molecular and phenotypic effects of different AHR genetic variants, we exploit a transgenic mouse model, termed “AHR-ratonized mice”, in which the endogenous AHR of C57BL/6 mice is ablated and either the rat wild-type (rWT), the deletion variant (DEL) or insertion variant (INS) is inserted into the mouse genome [26], as well as the TCDD-resistant DBA/2 mouse strain. The DEL isoform confers only moderate resistance to TCDD toxicity, relative to the INS isoform [26]. We compare hepatic transcriptomic responses of these transgenic mice to one another and to their corresponding TCDD-sensitive and resistant strains of rat. Results are then supplemented with whole-genome sequencing (WGS) to further isolate the specific genes responsible (*i.e.* gene “B”) for differential toxicities and to further characterize the mechanism by which TCDD activation of the AHR causes toxicity. These combined transcriptomic and genomic resources provide complimentary evidence for this and future studies.

## Results

### Experimental design

Our experimental strategy is outlined in Fig 1 and **S1 Table**. We examined TCDD-mediated transcriptional changes associated with various AHR isoforms within animals that have different backgrounds (Experiment #1, EXP1) or identical genetic backgrounds (Experiment #2, EXP2). Here we focus on liver as it shows large phenotypic differences between TCDD-resistant and TCDD-sensitive animals following exposure. Moreover, liver exhibits high expression of AHR [26], thereby making it an appropriate model for determining the effects of AHR isoforms on responses to TCDD. For EXP1, C57BL/6 and DBA/2 mice, along with rWT mice, were treated with a single dose of 0, 5 or 500 μg/kg TCDD in corn oil vehicle, with tissue collected 19 hours after. For EXP2, AHR-ratonized (INS/DEL/rWT) and C57BL/6 mice were treated with a single dose of 0, 125, 250, 500 or 1000 μg/kg TCDD in corn oil vehicle. These doses were selected so as to discriminate among the various strains/lines with regard to their overt toxicity responses. The approximate LD_50_ value for male C57BL/6 mice in our laboratory is 305 μg/kg [5], while male DBA/2 mice are reported to have a LD_50_ of 2570 μg/kg TCDD [4]. In the case of male rWT, DEL and INS mice, the dose of 500 μg/kg was lethal to 4/6, 2/6 and 0/6 animals respectively [26]. Thus, 5 μg/kg was definitely, and 125 μg/kg probably, sub-lethal to all animals in the present study, whereas the other doses would have been variably fatal over time. The first time point (19 hours) should reveal early (and thus most primary) changes in gene expression levels, while the second (4 days) is the time when histological alterations in the liver are first discernible in rats [27]. Hepatic tissue was collected 4 days following exposure and transcriptional profiling performed. Arrays were pre-processed independently for each group (**S1-3 Figs**). Transcriptomic data from 85 animals across 4 rat strains/lines were used for comparison [22,23]. Genes which demonstrated altered mRNA abundance as well as pathways showing significant enrichment for these genes only in the TCDD-sensitive (C57BL/6, rWT, L-E) or only in TCDD-resistant cohorts (INS, H/W) were identified. Finally for EXP3, WGS was performed on hepatic tissue gDNA extracted from untreated H/W and L-E rats. An average coverage of 85x and 15x was achieved for H/W and L-E respectively (Table 1). Single nucleotide variants (SNVs) were annotated with predicted impact and compared between rat strains (see **Methods**).

**Fig 1:**
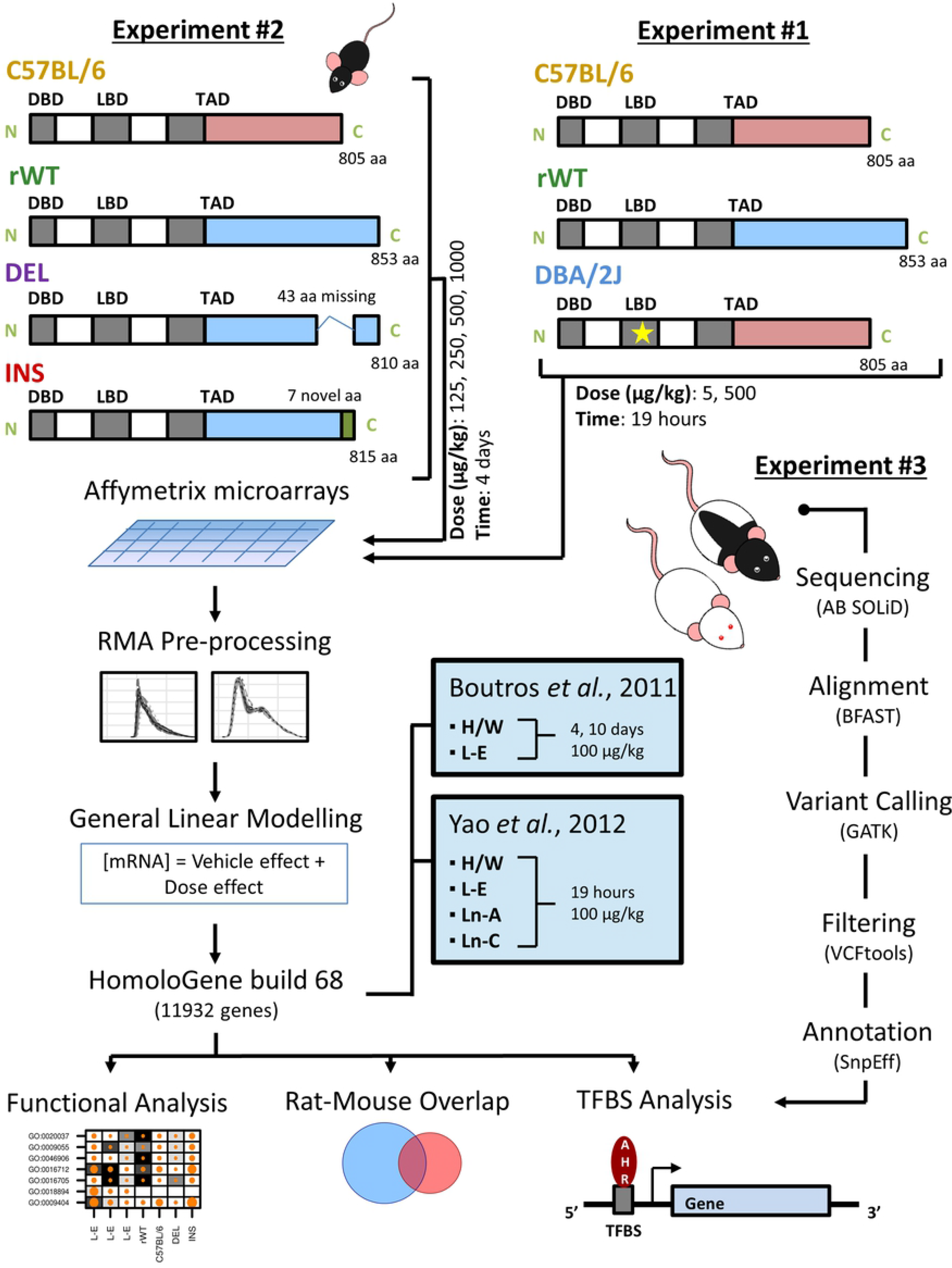
Experimental Design. This study evaluated hepatic transcriptomic profiles of 80 male mice that carry variant AHR isoforms (ratonized mice) – C57BL/6, rWT, DEL and INS – and treated with doses of 0, 125, 250, 500, or 1000 μg/kg TCDD in corn oil vehicle. Livers were excised at 4 days post-exposure and RNA abundance was profiled using microarrays. An additional 36 male mice (n = 12 each C57BL/6, rWT and DBA/2 (Ala375Val)) were treated with doses of 0, 5, or 500 μg/kg TCDD in corn oil vehicle and liver tissue collected 19 hours post-exposure. Data from two earlier studies that analyzed two strains of rats, Han/Wistar (H/W – INS and DEL) and Long-Evans (rWT), at various times post-treatment were included in the analysis, after filtering for orthologous genes using HomoloGene. Differentially abundant genes were subjected to pathway analysis using GOMiner, transcription factor binding site (TFBS) analysis and overlap visualization of significantly altered genes. Sequencing of genomic DNA isolated from liver of untreated L-E and H/W rats (n = 2 each) was performed and processed as shown. SNVs were identified and used to detect novel and/or lost TFBS within each rat strain. Genes containing such sites were further examined for changes to mRNA abundance using the above mentioned microarray data.

**Table 1:** Summary of L-E and H/W rat sequencing

| Strain | Average coverage | Number of Variants (pre-filter) | Number of Variants (post-filter) | Number of homozygous SNVs | Number of homozygous Indels | Number of “High Impact” Variants | Number of unique “High Impact” Variants |
| --- | --- | --- | --- | --- | --- | --- | --- |
| H/W | 85.34 | 3235191 | 1138926 | 176578 | 127870 | 756 | 35 SNVs, 608 indels |
| L-E | 14.73 | 2004271 | 1187745 | 161620 | 19520 | 220 | 116 SNVs, 14 indels |

Genomic DNA from hepatic tissue of H/W and L-E rats was sequenced using the AB SOLiD platform. Reads were aligned to the reference (rn6) using BFAST, followed by variant calling using GATK’s HaplotypeCaller. Variants were filtered to obtain only novel and unique high-quality variants for each strain, followed by annotation using SnpEff. Final numbers indicate total number of unique (H/W or L-E only, after removal of known variants [28]), homozygous, high impact variants.

### Transcriptomic responses to TCDD

TCDD treatment triggered changes in hepatic mRNA abundance in each of the mouse cohorts studied, but with substantial differences in the magnitude, direction and identity of genes affected. Animals treated with corn oil vehicle alone displayed similar transcriptomic profiles regardless of AHR isoform (adjusted Rand index (ARI) = 0.69 for TCDD, control) whereas TCDD-treated animals cluster more closely to animals with the same AHR isoform, regardless of dose (**S4A-B Fig**). Following linear modeling, a p-value sensitivity analysis was performed to compare different significance thresholds (**S4C-I Fig**). For downstream analyses, a dual threshold of effect-size (log_2_|fold change| > 1) and significance (p_adj_ < 0.05) was used to define transcripts with a statistically significantly difference in abundance following TCDD treatment.

### Early transcriptomic responses to TCDD differ by *Ahr* genotype

Previous studies observed transcriptomic changes as early as 6 hours in C57BL/6 mouse liver [29] and 19 hours in rat liver [21,23,30] following exposure to TCDD. To further study the role that the *Ahr* has in ‘early onset’ changes, we identified genes with significant differential mRNA abundance following a dose of 5 or 500 μg/kg TCDD in sensitive mouse strains (C57BL/6 or rWT) or resistant (DBA/2) mouse strains. We observed clear trends in response, with increased dose resulting in an increased number of differentially abundant transcripts (Fig 2A). The absolute number of changes to transcript abundance was considerably different between the groups at each dose tested. Intriguingly, liver from the ratonized rWT mouse demonstrated a heightened response relative to both the C57BL/6 and DBA/2 mice, even at the lowest dose tested. This may suggest differences in binding affinity for either TCDD and/or AHREs of the rWT *Ahr* relative to the mouse *Ahr*. As assessed by a modified sucrose gradient assay and Woolf plot, the apparent binding affinity of AHR for TCDD is quite similar in C57BL/6 mice and L-E rats harboring the WT receptor (Kd 1.8 *vs*. 2.2 nM respectively), while it is notably lower (16 nM) in the DBA/2 mouse strain [31,32]. However, after a lethal dose of TCDD, rWT mice tend to die much more rapidly compared with C57BL/6 mice [26], which probably bears on the difference in transcript abundance. In further support of this, at 500 μg/kg TCDD a similar number of differentially abundant transcripts was detected in C57BL/6 and DBA/2 mice, but in rWT mice, the number was still twice as high. We next examined the overlap of genes across these three groups (Fig 2B) and found 34 genes with differential mRNA abundance in all three groups. Five of these are part of the ‘AHR-core’ gene battery - a set of 11 well-documented TCDD-responsive genes with transcription previously shown to be mediated by the AHR [33,34,35,36] (Fig 2C). An additional four of these genes had differential RNA abundance in at least two groups: *Cyp1a2* in C57BL/6 and DBA/2 (borderline significant (log_2_ fold change = 0.99) in rWT) and *Fmo3*, *Nqo1* and *Ugt1a9* only in the TCDD-sensitive cohorts. *Inmt* shows significant repression in only rWT at this early time point, while *Aldh3a1* shows no response in mice (consistent with previous studies [29,37,38]). We next examined sets of genes which demonstrated altered RNA abundance in the TCDD-resistant DBA/2 mouse liver or TCDD-sensitive cohorts (Fig 2D-E). Genes with altered mRNA abundance in DBA/2 mice included *Apol7c*, *Tnfaip8l3* and *Htatip2* (both low and high dose exposure). These genes were similarly altered in the sensitive strains. *Rpl18a* and *Mbd6* had altered mRNA abundance exclusively in the resistant group (high and low dose), while two additional genes, *Onecut2* and *Lipg* had altered mRNA abundance exclusively in the resistant mouse, high dose group (Fig 2D). Alternatively, Fig 2E highlights differentially abundant transcripts that appear exclusively in sensitive animals, regardless of dose (including *Acpp*, *Dclk3*, *Fmo2*, *Pmm1* and *Ugdh*) and genes that respond in only sensitive strains (such as *Acot2*, *Acot3* and *Smcp*). Pathway analysis suggested that genes with altered RNA abundance in DBA/2 mice are involved in lipase activity, while the sensitive strains both demonstrate an enrichment of genes involved in xenobiotic and flavonoid metabolic processes as well as myristoyl-and palmitoyl-CoA hydrolase and oxidoreductase activities.

**Fig 2:**
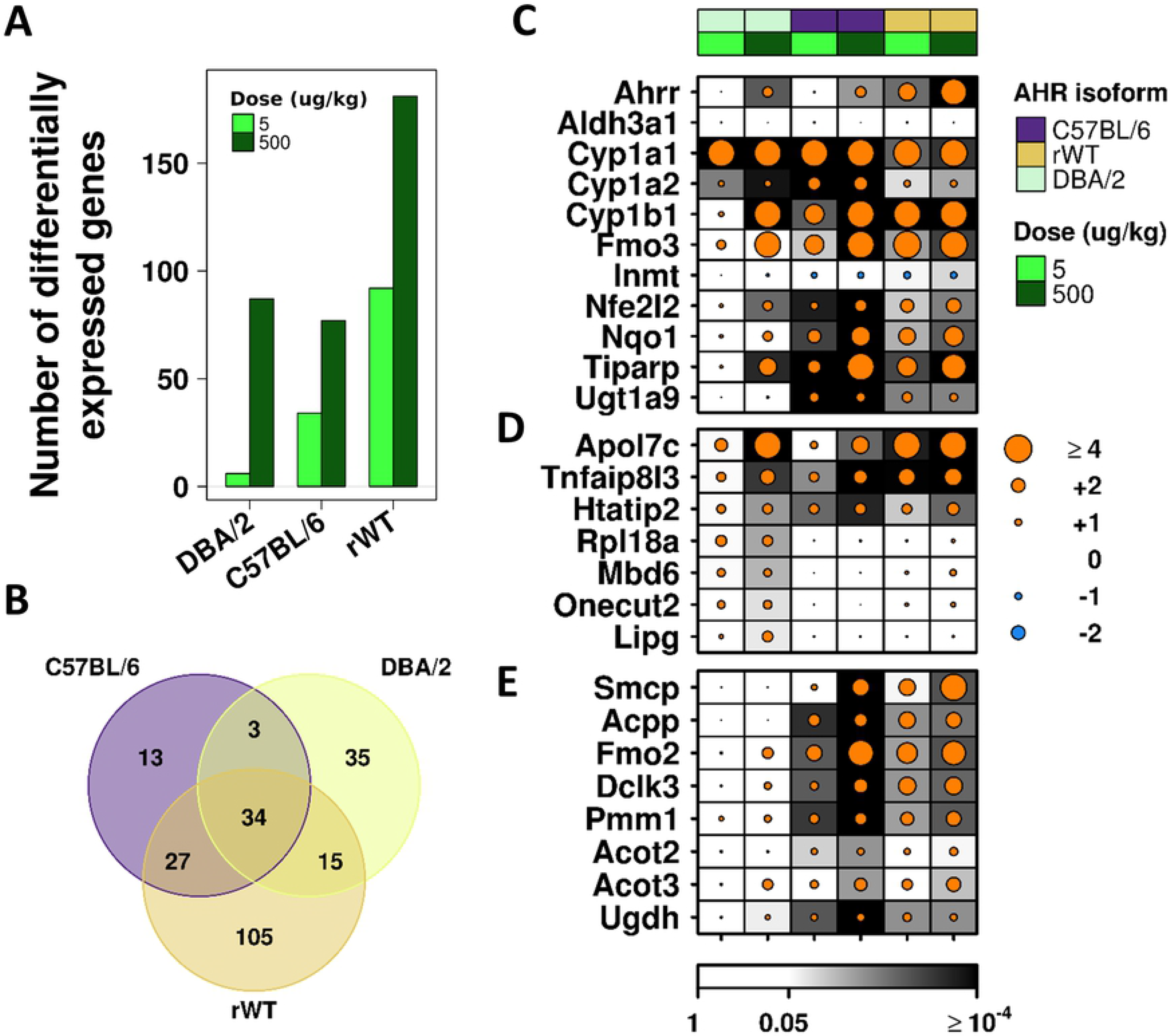
Differential transcriptomic profiles emerge early following exposure to TCDD. (A) Using a dual threshold of |log_2_ fold change| > 1, p_adj_ < 0.05, genes with differential mRNA abundance were identified. As expected, the TCDD-resistant DBA/2 mouse liver showed a transcriptional response following only the high dose of TCDD, while the sensitive C57BL/6 and rWT strains demonstrated considerable changes following low exposure that increased with dose. (B) Overlap of these genes in each cohort, following exposure to 500 μg/kg TCDD for 19 hours. Fold change of (C) “AhR-core” genes, (D) genes with significantly altered mRNA abundance in resistant mouse liver or (E) sensitive strains. Dot size indicates magnitude of change following exposure to TCDD relative to controls, while colour indicates direction of change (orange = increased abundance, blue = decreased abundance); background shading indicates FDR-adjusted p-value.

### Late transcriptomic responses to TCDD in ratonized mice

We next sought to identify changes in transcriptomic patterns that occur late following exposure to TCDD. Four days after exposure to TCDD, a considerable difference in the hepatic transcriptomic profiles of H/W and L-E rats has been observed [22]. Therefore, we evaluated the responses of ratonized transgenic mice at this same time point and utilizing a dose-response experiment. As above, we identified a large number of genes with differential mRNA abundance in the TCDD-sensitive rWT mouse, as well as the more resistant DEL mouse, and a muted response in highly resistant INS mouse (Fig 3A). As a clear dose-response regarding number of differentially abundant RNAs was not apparent, we focused downstream analyses to the 500μg/kg TCDD group, for consistency with EXP1. As explained above, this dose is above the LD_50_ for the TCDD-sensitive mice (C57BL/6 and rWT) but below it in TCDD-resistant cohorts (DEL and INS; [26]). Interestingly, the largest overlap occurred between the DEL and rWT isoforms, consistent across all doses (Fig 3B, **S5B-D Fig**) and consistent with a previous study demonstrating reduced protection against TCDD toxicity by this DEL isoform in these models [26]. A total of 15 genes were identified in all four groups, including six of the eleven “AHR-core” genes (Fig 3C). As expected, responses to TCDD of *Cyp1a1* and *Cyp1b1* were consistent across all AHR isoforms and all doses [26]. Similarly, *Ahrr*, *Nqo1*, *Tiparp* and *Ugt1a9* showed consistent changes at the 500 μg/kg TCDD dose across all strains. Interestingly, *Inmt* did not show changes to mRNA abundance in the C57BL/6 or INS but did show dramatic repression in the DEL and rWT, indicating a response specific to these AHR-genotypes. Exclusive to the TCDD-sensitive cohorts, 28 genes were identified with altered transcriptomic response to TCDD, with only three genes exclusive to the TCDD-resistant cohorts. Interestingly, this included *Fmo2* (Fig 3D, **right panel**) which was altered exclusively in the TCDD-sensitive cohorts at the early time point used in EXP1 (Fig 3D, **left panel**) suggesting a short-lived induction in sensitive strains/lines that is delayed in resistant ones. It is noteworthy, though, that the same resistant models were not used in both time-points, and thus *Fmo2* induction may have had a decreasing trend also in DEL and INS mice over time.

**Fig 3:**
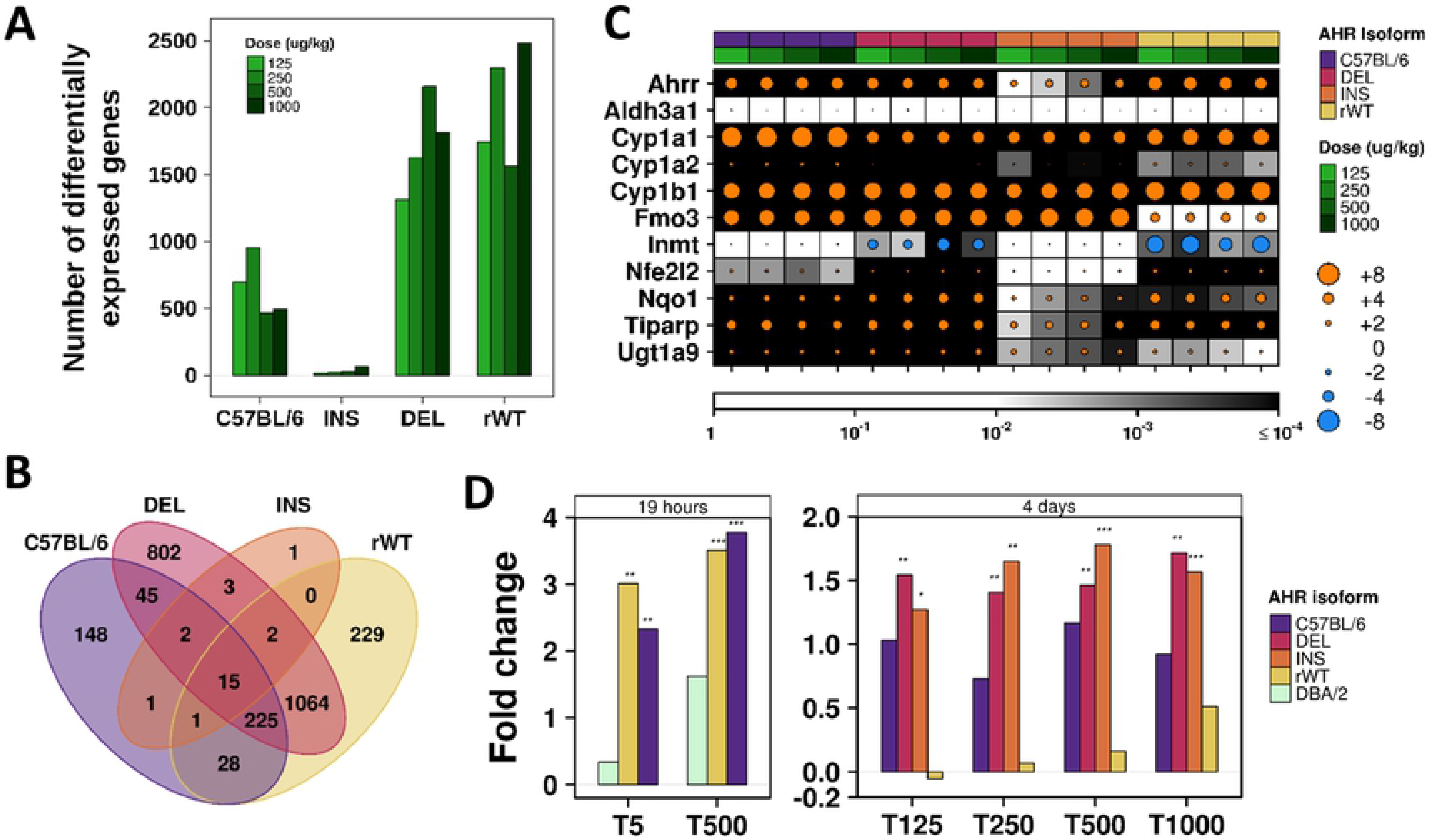
Late transcriptomic changes in AHR ratonized mouse liver. (A) Using a dual threshold of |log_2_ fold change| > 1, p_adj_ < 0.05, genes with differential mRNA abundance were identified for each AHR isoform. A clear dose-response pattern was not observed, however a considerable increase in the number of genes demonstrating altered mRNA abundance was detected between the TCDD-resistant INS isoform and TCDD-sensitive rWT and C57BL/6 isoforms, and less sensitive DEL isoform. (B) Overlap of these genes in each cohort, following exposure to 500 μg/kg TCDD for 96 hours. (C) Fold change of “AhR-core” genes ordered by AHR isoform and increasing exposure. Dot size indicates magnitude of change following exposure to TCDD relative to controls (log_2_ fold change), while colour indicates direction of change (orange = increased abundance, blue = decreased abundance); background shading indicates FDR-adjusted p-value. (D) *Fmo2* demonstrated significant changes in mRNA abundance earlier among TCDD-sensitive strains that appear later in TCDD-resistant ones. Bar height shows magnitude of change (log_2_ fold change); * p < 0.05, ** p < 0.01, *** p < 0.001.

### Differences between rat and mouse hepatic response to TCDD

We next expanded the study by contrasting our findings with a rat-transcriptomic dataset previously generated under similar experimental conditions [22,23,30]. We evaluated 11,932 orthologous genes, obtained from EXP1 and EXP2 (500 μg/kg TCDD), L-E and H/W rats (1/4/10 days, 100 μg/kg TCDD) and Ln-A and Ln-C rats (19 hours, 100 μg/kg TCDD). Using the same dual threshold of |log_2_ fold change| > 1, p_adj_ < 0.05, we found little overlap between species (with a higher degree of overlap among different strains/lines of the same species regardless of TCDD response phenotype; Fig 4). Thus, in the transgenic mice, the host species was a more important determinant of the resultant responsiveness than the AHR isoform. Four ‘AHR-core’ genes showed altered mRNA abundance in all cohorts (*Cyp1a1*, *Cyp1b1*, *Nqo1* and *Tiparp*). *Nfe2l2* showed altered transcript abundance in all cohorts except INS from EXP2 and *Inmt* was repressed more consistently in rats (H/W and L-E liver at 19 hours, 4 and 10 days, and Ln-A and Ln-C at 19 hours) than mice (near significant in rWT at 19 hours, DEL/rWT at 4 days, all doses).

**Fig 4:**
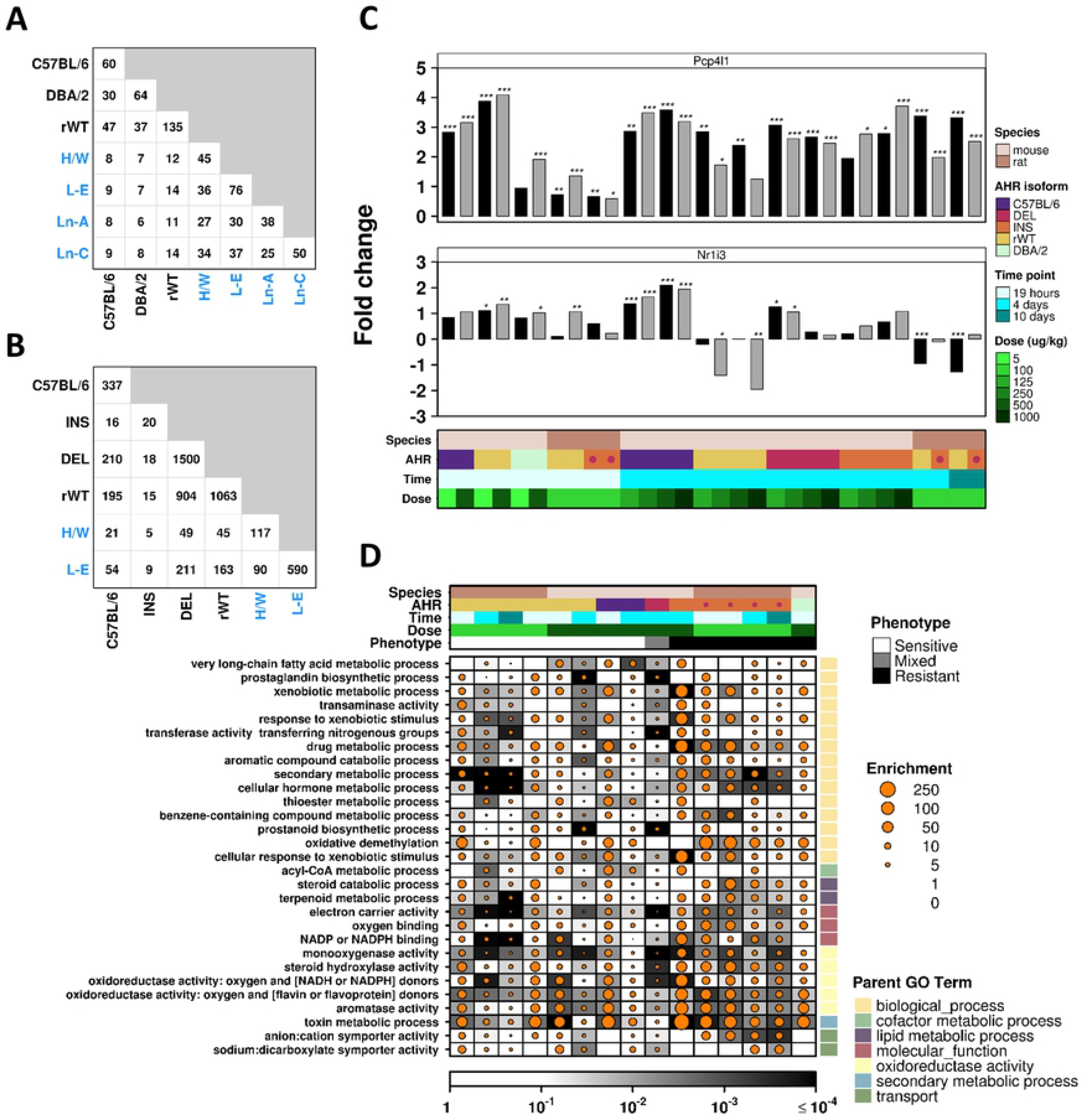
Comparison of transcriptomic changes between species and AHR isoform. Intersection of the number of genes with significantly altered mRNA abundance (|log_2_ fold change| > 1, p_adj_ < 0.05) in each examined cohort. Responses following treatment with 500 μg/kg (mouse strains) or 100 μg/kg (rat strains, blue text) TCDD for (A) 19 hours or (B) 4 days. Only orthologous genes were examined. (C) *Pcp4l1* demonstrated significant changes in mRNA abundance among most cohorts. Bar height shows magnitude of change (log_2_ fold change, all with increased abundance relative to controls); * p < 0.05, ** p < 0.01, *** p < 0.001. (D) Pathway analysis of significantly differentially abundant orthologous genes in mouse and rat cohorts was performed using GoMiner. Significantly enriched biological pathways (p_ad*j*_ ≤ 0.01, enrichment > 15) were identified within each group and status is shown across all groups. Dot size indicates enrichment score while background shading represents significance level. Empty cells indicate 0 genes within that pathway were differentially abundant.

Of the 28 genes identified above in the sensitive cohorts (EXP2), 18 had homologs present in the rat dataset. Of these, *Igfbp3* demonstrated differential mRNA abundance (|log_2_ fold change| > 1, p_adj_ < 0.05) in both the TCDD-sensitive L-E rats and TCDD-resistant H/W rats following a 4-day exposure while *Pxdc1* was altered only in L-E rat liver and *Cyp1a2* was altered only in H/W rat liver (though this showed near significant induction in all animals tested). PX domain-containing protein 1 (*Pxdc1*) is significantly repressed in TCDD-sensitive cohorts (4 days following exposure, regardless of dose in C57BL/6, rWT, as well as in L-E rats; near significant in DEL mice). This gene is poorly characterized and differential transcript abundance could not be directly attributed to the AHR because this gene demonstrated altered presence of Transcription Factor Binding Sites (TFBSs) among the species and strains/lines used (**S2 Table**). Additionally, no genomic differences were detected between the TCDD-sensitive L-E and TCDD-resistant H/W rats (**S3 Table**), suggesting that *Pxdc1* is a poor candidate for the proposed gene “B”.

*Pcp4l1* demonstrated induced RNA abundance in most cohorts (Fig 4C, **top**). Genomic analyses indicated the presence of multiple AHRE-1 (core, extended) and ARE motifs within the promoter region of this gene in both species; however, with more occurrences in mice (n = 8 AHRE-1 (core) and 2 ARE motifs) than rats (n = 7 AHRE-1 (core) and 1 ARE motifs). *Pcp4l1* encodes Purkinje cell protein 4-like 1 and is typically expressed in neuronal tissue; it has been hypothesized to be a calmodulin inhibitor [39]. Interestingly, this gene is adjacent to and in the reverse orientation of, *Nr1i3* – a gene encoding a transcription factor previously associated with enhanced TCDD sensitivity [29], showing increased mRNA abundance in TCDD-sensitive male mice than TCDD-resistant female mice. *Nr1i3* demonstrates altered transcriptional response following TCDD exposure across species (Fig 4C, **bottom**): hepatic mRNA abundance was increased after TCDD exposure in C57BL/6 mice (4 days, all doses) and earlier in rWT mice and L-E rats (19 hours). This response was followed by a significantly reduced mRNA abundance in rWT mice (4 days) and L-E rats (4 and 10 days). This gene also shows different presence of AHREs in its promoter region between species (n = 2 AHRE-1 (core) motif in both mice and rats; 4 ARE motifs in mice; 1 AHRE-2 and 1 ARE motif in rats). No differences in AHREs were observed between H/W and L-E rats for either *Pcp4l1* or *Nr1i3*. This makes these genes interesting candidates for involvement in TCDD-induced toxicity.

Finally, functional pathways affected by TCDD were compared across these datasets, using only orthologous genes (**S4 Table**). Unsurprisingly, sensitive strains exhibit a larger number of significantly enriched pathways than do resistant animals, many of which are altered in multiple strains/lines (Fig 4D). Specifically, TCDD-sensitive animals display a more significant enrichment of altered transcripts among metabolic and oxidoreductase activity pathways than resistant animals whereas resistant animals exhibit responses mostly in metabolism-related processes and transport activities.

### Identifying candidates for gene “B” using genomic variants

The H/W rat strain has astonishing resistance to TCDD toxicities, predominantly conferred by a point mutation in the transactivation domain of the *Ahr* gene [18]. Part of this resistance has also been contributed to a second hypothesized gene, termed gene “B” [24]. Of the 642 nuclear “high impact” homozygous variants unique to the H/W strain (**S3 Table**), 209 mapped to genes evaluated in our microarray cohorts. Twenty of these did not show altered transcript abundance in any of our cohorts. Five genes demonstrated altered mRNA abundance exclusively in H/W rat (**S6 Fig**), and 125 showed TCDD-mediated mRNA abundance changes in at least one experimental group, excluding H/W rat liver (28 of these showed significantly altered mRNA abundance in at least 8 non-H/W groups). None of these gene-sets contained more variant genes with altered mRNA abundance than expected by chance alone (hypergeometric test, p > 0.05). Additionally, a single “high impact” homozygous mitochondrial SNP was identified in H/W that results in a lost stop codon in mitochondrial gene NADH dehydrogenase 6 (*MT-nd6*). This was not found in other rat strains.

Since the above list does not include variants located within intergenic regions, an alternative analysis was performed to identify genes demonstrating altered transcript abundance associated with modified regulatory regions. Specifically, we searched the H/W genome for novel or lost transcription factor binding sites (TFBSs) specific to the AHR with the hypothesis that a gain of a TFBS may allow AHR-mediated transcription of a TCDD-resistance gene whereas a loss of a TFBS would prevent transcription of a TCDD-susceptibility gene. In total, 13.4% of genes (1,446 of 10,772 with available TFBS data) had either a gain or loss of at least one of the AHREs examined (**S2 Table**). Of these, 32 exhibited changes to the number of AHRE-1 (full) motifs within their promoter regions as compared to the TCDD-sensitive L-E rat (Table 2), with 17 showing a gain and 15 a loss. Perhaps more interestingly, 15 genes harboured this motif in H/W (with none identified in the same region for L-E), while nine genes demonstrated a complete loss of this motif from the promoter region in H/W rats. TCDD-responsive genes were not enriched in those demonstrating a novel AHRE-1 (full) motif in H/W and TCDD-responsiveness in H/W liver (hypergeometric test; **S5 Table**).

We next evaluated differential transcriptional patterns for genes demonstrating differences among these TFBSs between H/W and L-E rats. For example, *Cxxc5* shows a loss of the AHRE1 (full) motif within its promoter region in H/W (one found in H/W and two in L-E/rn6) and exhibits significantly reduced transcript abundance in liver from both L-E (4 and 10 days) and the TCDD-sensitive Ln-C rat (19 hours) following exposure to TCDD (Table 2). Similarly, *Sugp1* demonstrated a complete loss of this motif from the H/W strain relative to L-E (n = 0 and 1 respectively), with mice (mm9) similarly lacking the AHRE-1 (full) motif. This gene shows significantly increased mRNA abundance in only the TCDD-sensitive L-E rat liver (4 day, log_2_ fold change = 0.33, p_adj_ = 0.0012). Alternatively, *Hgfac* demonstrates a novel AHRE-1 (full) motif in the H/W rat (n = 1 in H/W and 0 in L-E/rn6) and its’ mRNA abundance is significantly increased in both strains, 4 and 10 days following treatment, but is much higher in L-E rat liver (Table 2). No changes were detected in the AHR-ratonized mice, and this gene was not included on the arrays used for 1 day exposures, preventing assessment in the Ln-A and Ln-C rats. This list supplies a robust set of 12 gene “B” candidates, in particular *Cxxc5*, with evidence of altered TFBSs and altered transcriptional patterns in only rats that may prove suitable for further mechanistic investigation.

**Table 2:**
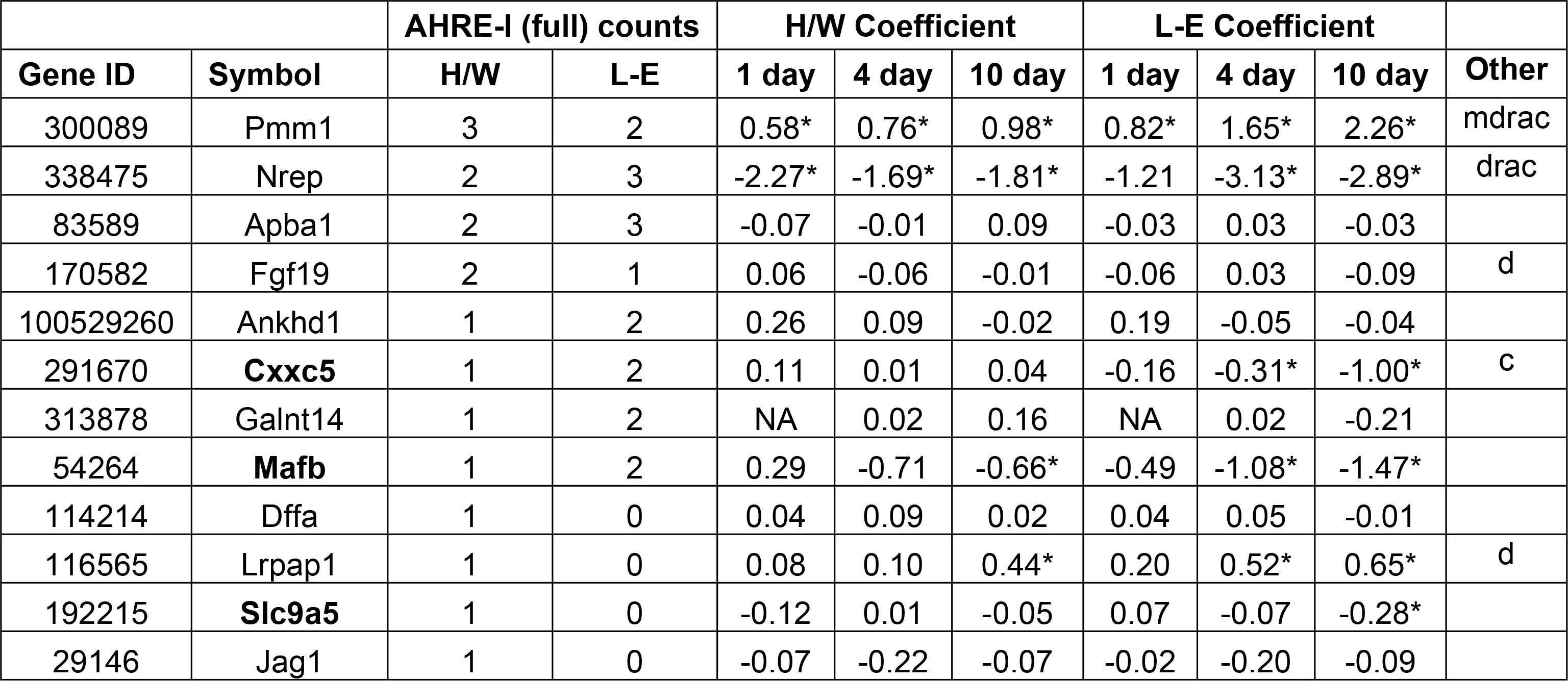
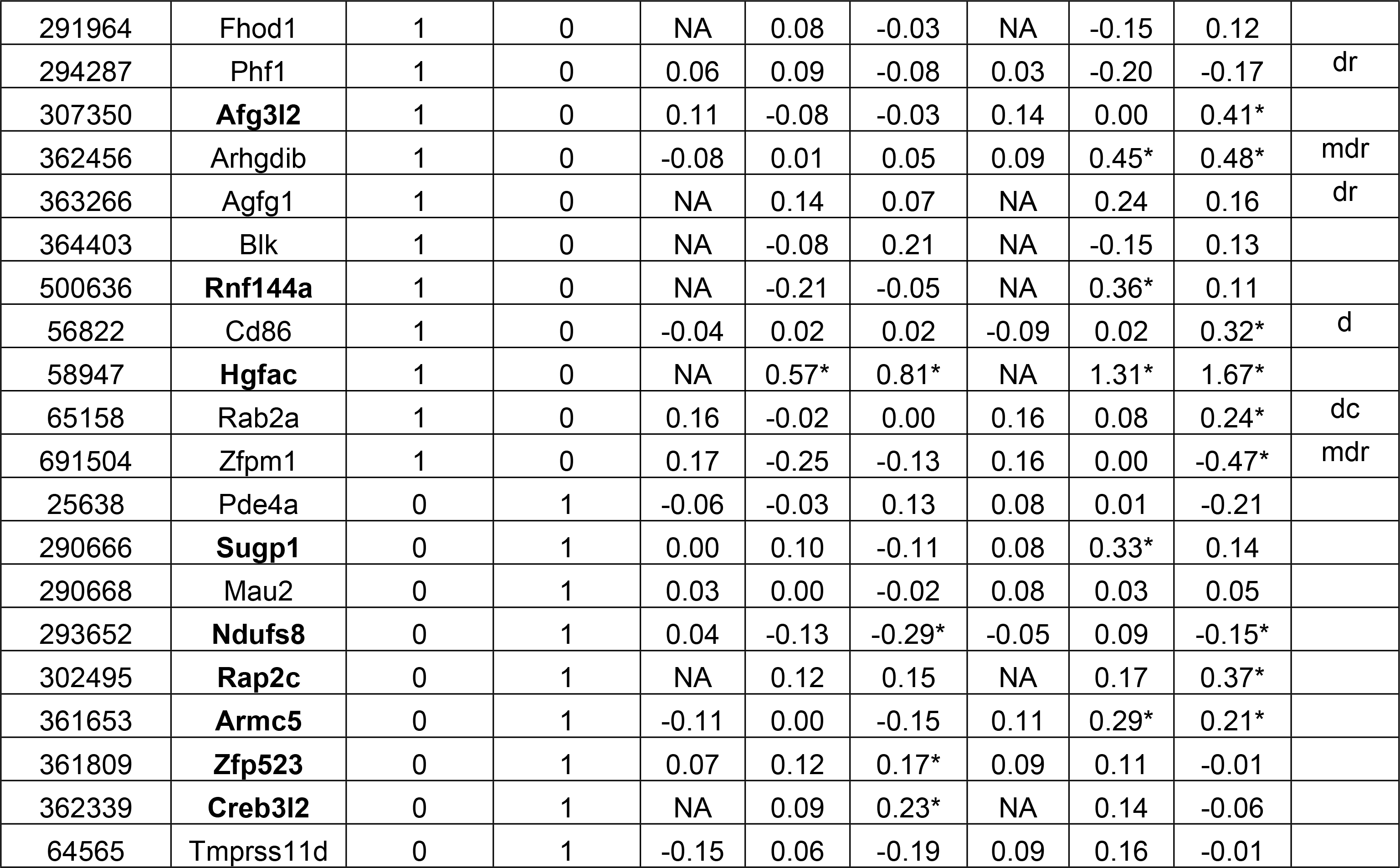
Genes demonstrating altered transcription factor binding sites

The number of occurrences for the AHRE-I (full) motif within a ±3 kbp region around the transcription start site for each gene was determined for each H/W and L-E rat. Genes which demonstrate either a gain or loss of this motif in H/W relative to L-E may represent gene “B”. Of the 11,392 rat/mouse orthologous genes examined, 32 were revealed to have either a gain or loss of this motif in H/W rats. Of these, 19 showed altered mRNA abundance following treatment with TCDD in at least one rat strain at one or more time points evaluated. *significantly altered mRNA abundance in TCDD treated group relative to controls (p_adj_ < 0.05). Column labeled ‘Other’ indicates significantly altered mRNA abundance in: a = Ln-A; c = Ln-C (100 μg/kg TCDD, p_adj_ < 0.05); m = C57BL/6 mouse; d = DEL, i = INS, r = rWT AhR-ratonized mice (500 μg/kg TCDD, p_adj_ < 0.05). Gene symbols highlighted with bold font indicate key gene “B” candidates (n = 12).

## Discussion

There is abundant evidence that the AHR structure is a primary determinant of susceptibility to TCDD toxicity [12,13,14,15,16,19,reviewed in 40]. Two rodent species, mice and rats, show inter-strain differences in response to TCDD-insult depending on AHR genotype [22,23,41]. Furthermore, previous studies have shown little overlap in the transcriptomic response following TCDD exposure in TCDD-sensitive strains of mouse *vs*. rat [20,21]. These differences may in part be due to the intrinsic genetic variation between model organisms. In order to remove this confounding factor, we examined transgenic mouse models harbouring different rat AHR isoforms within an identical genetic background. To further support this analysis, we performed whole-genome sequencing of two common rat models with highly different responses to TCDD exposure, due to bearing the variant AHR isoforms utilized in our transgenic models. By contrasting the profiles of homozygous SNVs in each strain with the transcriptional landscape of various rat and mouse models, we aim to identify phenotype pertinent genes.

Previous studies contrasting basal transcriptomic profiles of mice and rats with different AHR genotypes [15,42] found large differences in mRNA abundance of many genes in the absence of xenobiotic AHR ligands. Contrary to this, there was little difference among basal transcriptomic profiles among our transgenic mice, suggesting that these variations in AHR structure have little effect on basal gene expression. This may be partly explained by differences in the mRNA abundance of *Ahr* itself – in liver tissue, basal expression of TCDD-sensitive *Ahr* isoforms (rWT, DEL) is considerably higher than for the TCDD-resistant INS isoform in these transgenic mice [26]. This represents an intriguing pattern and will require further study to better understand the inherent differences in activity of these isoforms.

We found that the AHR alone is sufficient to explain much of the observed variation in sensitivity to TCDD. At all doses assessed, roughly 50% of genes demonstrating TCDD-mediated changes in transcription were unique to a single transgenic model. For example, at the highest dose studied (1000 μg/kg TCDD), 33.8%, 11.7%, 5.2%, and 0.3% of differentially abundant transcripts were specific to the rWT, DEL, C57BL/6 and INS cohorts respectively (of a total 3,076 with altered mRNA abundance). These proportions were fairly consistent across doses, with the proportion of genes with altered mRNA abundance that were unique to C57BL/6 negatively correlated with dose, while INS showed a positive correlation; DEL and rWT were less consistent, with each showing ~30% unique changes at 500 or 1000 μg/kg TCDD respectively. At first glance, it seems unusual that a considerable transcriptomic response was detected in the transgenic model expressing the TCDD-semi-resistant H/W DEL isoform. Both the DEL and INS isoforms are expressed naturally in the H/W rat, in roughly 15% to 85% proportions, however the DEL isoform displays a significantly higher intrinsic transactivation activity [19], partially explaining the reduced resistance observed in these models [26]. Since our ratonized mice varied only at the AHR locus, the remaining 50% of differentially transcribed genes may be due to additional transcription factors as either primary or secondary effects, as suggested by the low overlap between early (EXP1) and late (EXP2) studies.

We identified twelve candidates for the hypothesized TCDD-resistance associated gene “B”. These genes demonstrate altered TFBS within their promoter regions and altered transcript abundance in rats, but not the corresponding AHR-ratonized mice. Here we focused on the AHRE-1 (full) motif, because it results in the most productive receptor-DNA interaction [43]. In particular, three genes were identified that had lost this motif in H/W rats, and demonstrated altered mRNA abundance in only L-E (*Sugp1*, *Rap2c* and *Armc5*). SURP and G patch domain containing 1 (*Sugp1*) encodes a splicing factor that had increased mRNA abundance in TCDD-exposed L-E rat liver (4 days); however the magnitude of change was small (log_2_ fold change = 0.33). *Rap2c* is member of RAS oncogene family that showed increased mRNA abundance in TCDD-exposed L-E rat liver (10 days), again however, the magnitude of change was small (log_2_ fold change = 0.37). Armadillo repeat containing 5 (*Armc5*), showed significantly increased mRNA abundance following treatment in L-E rat liver, as well as in liver of mice expressing the rWT-*Ahr*, while it was repressed in livers of C57BL/6 mice (despite also containing a proximal AHRE-1 (full) motif). Little is known about this gene; however, it is a putative tumour suppressor gene [44].

Genomic sequencing further revealed loss of one of two AHRE-1 (full) occurrences from the promoter region of the CXXC finger protein 5 (*Cxxc5*) in the H/W rat along with significantly repressed mRNA abundance in livers of TCDD-sensitive L-E (10 days, log_2_ fold change = −1, p_adj_ = 3.1×10^−6^) and Ln-C (19 hours, log_2_ fold change = −0.45, p_adj_ = 0.014) rats. Of interest, the transcription factor CXXC5 has been shown to inhibit expression of cytochrome c oxidase by binding to an oxygen response element in the proximal promoter of *Cox4l2* in human lung cells [45]. The delayed onset of repression of *Cxxc5* in TCDD-sensitive cohorts may be a secondary response due to the presence of oxidative stress, and possibly corresponds with altered energy metabolism observed in these animals. Another gene, *Mafb* (v-maf musculoaponeurotic fibrosarcoma oncogene family, protein B), has a single AHRE-1 (full) motif in H/W rats but two in L-E rats and exhibited significantly repressed mRNA abundance in L-E rats. This may bear on the disparate developmental toxicity outcomes upon TCDD exposure observed in these strains: cleft palate was not seen at any dose in H/W rat progeny but it occurred in 71.4% of offspring in L-E rats treated with 5 μg/kg TCDD [46]. In humans, *Mafb* variants have been associated with cleft palate and lip [47]. Finally, a novel AHRE-1 (full) motif was detected in the promoter region of *Hgfac* (HGF activator) in H/W rats that was absent in L-E rats. mRNA abundance for this gene was significantly increased in TCDD-exposed liver from L-E rats (4 and 10 days, log_2_ fold change = 1.3 and 1.7, p_adj_ = 1.16×10^−12^ and 1.16×10^−17^) with a more muted response in H/W rats (4 and 10 days, log_2_ fold change = 0.57 and 0.81, p_adj_ = 3.7×10^−4^ and 4.28×10^−8^). These genes provide interesting candidates for gene “B” that require further studies into its potential involvement in the onset of TCDD-toxicities.

The purpose of this study was to ascertain the mechanism of classic TCDD toxicity using various model systems, including transgenic mice to compare various rat *Ahr* variants in a system with a homogeneous genetic background, and various strains of rat, each with differing phenotypic responses to TCDD. To accomplish this, we generated unique transcriptomic and genomic datasets that provide multiple levels of evidence. Using this valuable resource, we identified several genes whose transcription was selectively altered by TCDD in either TCDD-sensitive or TCDD-resistant cohorts, a differential response that can be attributed to the particular AHR isoform expressed in each cohort. *Pxdc1* in particular demonstrated differential transcription between TCDD-sensitive and TCDD-resistant models across both mice and rats. However, the transcriptional responsiveness of this gene could not be explained by genomic differences in AHR-binding sites, as the transcription factor binding site analysis revealed highly variant sites between species, and no major difference between strains of rat. However, genomic sequence analysis allowed identification of differences between sensitive and resistant rat strains, which are potential “gene B” candidates. For instance, *Cxxc5* was found to have fewer occurrences of AHRE-1 (full) TFBSs in H/W relative to L-E, and had reduced RNA abundance in sensitive strains. This is a suitable candidate for further study in relation to mechanisms of TCDD toxicity and regulatory roles of the AHR.

## Materials and Methods

### Animal handling

Three separate experiments were performed (Fig 1). In the first, adult male C57BL/6*Kuo*, rWT and DBA/2J mice were evaluated. In the second, adult male C57BL/6 mice carrying 4 different *Ahr* isoforms were examined: C57BL/6*Kuo* and rWT, DEL, and INS transgenic mice. Transgenic mice were generated as described previously [26]. Briefly, animals were bred from *Ahr*-null mice to avoid interference by the C57BL/6 *Ahr*. Once established, transgenic colonies were bred at the National Public Health Institute, Division of Environmental Health in Kuopio, Finland. For this study, animal ages varied from 12–23 weeks. All animals were housed singly in Makrolon cages. The housing environment was maintained at 21 ± 1°C with relative humidity at 50 ± 10%, and followed a 12-hour light cycle. Tap water and R36 pellet feed (Lactamin, Stockholm, Sweden) or Altromin 1314 pellet feed (DBA/2 mice; Altromin Spezialfutter GmbH & Co. KG, Lage, Germany) were available *ad libitum.*

In the first experiment (EXP1), a total of 36 mice were used (n = 12 per genotype), divided into groups of four mice per treatment group. Mice were treated by oral gavage with a single dose of 0, 5, or 500 μg/kg TCDD dissolved in corn oil vehicle (**S1 Table**). Animals were euthanized by carbon dioxide, followed immediately by cardiac exsanguination 19 hours following treatment. In the second experiment (EXP2), a total of 84 mice were used, divided into groups according to *Ahr* isoform. Mice were treated by oral gavage with a single dose of 0, 125, 250, 500, or 1000 μg/kg TCDD dissolved in corn oil vehicle (**S1 Table**). Animals were euthanized by cervical dislocation 4 days following exposure and their livers excised. A single rWT animal from the 1000 μg/kg TCDD group died prematurely and was thus excluded from the study, thereby leaving 83 animals.

All study plans were approved by the Finnish National Animal Experiment Board (Eläinkoelautakunta, ELLA; permit code: ESLH-2008-07223/Ym-23). All animal handling and reporting comply with ARRIVE guidelines [48].

### Microarray hybridization

Mouse livers were frozen in liquid nitrogen immediately upon excision and stored at −80°C. Tissue samples were homogenized and RNA was isolated using an RNeasy Mini Kit (Qiagen, Mississauga, Canada) according to the manufacturer's instructions. Total RNA was quantitated using a NanoDrop UV spectrophotometer (Thermo Scientific, Mississauga, ON) and RNA quality was verified using RNA 6000 Nano kits on an Agilent 2100 Bioanalyzer (Agilent Technologies, Mississauga, ON). RNA abundance levels were assayed using Affymetrix Mouse Gene 1.1 ST arrays (Affymetrix Mouse Gene 2.0 ST arrays were used for the C57BL/6 mice for EXP1) at The Centre for Applied Genomics (TCAG; Toronto, ON) as described previously [22].

### Data pre-processing and statistical analysis

Raw data (CEL files) were loaded into the R statistical environment (v3.4.3) using the affy package (v1.48.0) [49] of the BioConductor library [50] and preprocessed using the RMA algorithm [51]. Each group (experiment/AHR isoform) was preprocessed independently to avoid masking any differences between isoforms. Probes were mapped to EntrezGene IDs using the custom mogene11stmmentrezgcdf and mogene20stmmentrezgcdf packages (v22.0.0) [52]. Visual inspection of array distribution and homogeneity of the results suggested the presence of outliers. These were further identified using Dixon’s Q test [53] as implemented in the outliers package (v0.14) in R, which was used to compare abundance patterns of “AHR-core” genes for each genotype/treatment combination. Three TCDD-treated rWT animals were removed from EXP2, along with one control DBA animal and one TCDD-treated rWT animal from EXP1, as mRNA abundance for these AHR-core genes closely resembled the control animals (FDR-adjusted Dixon’s Q test, *p* < 0.05 for 2 or more AHR-core genes, requiring at least one of *Cyp1a1*, *Cyp1a2* or *Cyp1b1*; **S1 Fig**). Remaining arrays were re-processed without outliers (**S2 and S3 Figs**). ComBat was run on the combined dataset, using the sva package (v3.24.4) for R, to adjust RMA normalized values for comparison across batches (**S4A Fig**). General linear modeling was performed to determine which genes were significantly altered following exposure to TCDD. Specifically, for each gene (i) the abundance (Y) was modeled as Y_i_ = β_0_ + β_1_X, where X indicates the dose of TCDD administered. Each experiment/AHR isoform was modeled independently, using RMA normalized values (without ComBat correction as each group was processed as a single batch) with each dose treated as a factor. Genes were modeled as a univariate combination of a basal effect (β_0_) (represented by the vehicle control) and TCDD effect (β_1_) with contrasts fit to compare each dose-effect to baseline. Standard errors of the coefficients were smoothed using an empirical Bayes method [54] and significance was identified using model-based t-tests, followed by FDR adjustment for multiple testing [55]. Sensitivity of results to various p_adj_-value cut-offs was assessed (Fig 2B, **S4C-I**). Those genes deemed significantly altered (p_adj_ < 0.05 and |log_2_ fold change| > 1) following treatment were examined in downstream analyses. All statistical analyses were performed using the limma package for R (v3.32.10) [54]. Venn diagrams were created using the VennDiagram package (v1.6.21) for R [56] to visualize overlap between groups. All other data visualizations were generated using the BPG plotting package (v5.9.8) [57], using the lattice (v0.20-38) and latticeExtra (v0.6-28) packages for R.

### Rat-mouse overlap analysis

Publicly available data for male Han/Wistar (H/W) and Long-Evans (L-E) rats, as well as two lines (Ln-A and Ln-C) derived from L-E × H/W crosses, were used for comparison [22,23,30,38]. Data consisted of the hepatic transcriptomic responses of rats to a single dose of TCDD (100 μg/kg in corn oil) at three time points (19 hours, 4 and 10 days). All data were available from the TCDD.Transcriptomics (v2.2.5) package [38] for R. Due to the use of different microarrays between studies, HomoloGene (build 68) was used to identify orthologous genes. Homologene IDs which mapped to multiple EntrezGeneIDs were discarded. In total, 11,932 genes were found to be orthologous between species and were used for overlap analyses (**S6 Table**).

### Pathway analysis

Pathway analysis was conducted in order to elucidate the functions of genes demonstrating significantly altered mRNA abundance for each AHR isoform cohort. Analysis was performed using the High-Throughput GoMiner web interface (application build 469, database build 2011-01, accessed 2019-02) [58]. A separate run was performed for each mouse AHR isoform and each available rat dataset. We compared significantly altered genes against a randomly drawn sample from all orthologous mouse (or rat) genes in the dataset, using an FDR threshold of 0.1, 1000 randomizations, all mouse (or rat) databases and look-up options, and all GO evidence codes and ontologies (**S4 Table**). Results were further filtered using a threshold of p_adj_ < 0.01 and an enrichment score > 15 in at least one of the examined groups. This resulted in 29 unique gene ontologies to examine further.

### Genome sequencing

Untreated adult male outbred H/W (*Kuopio*) and inbred L-E (*Turku/AB*) [59] rats were euthanized by decapitation and their livers excised. Tissue was frozen in liquid nitrogen and shipped to the analytic facility on dry ice. Genomic DNA (gDNA) was isolated and whole genome sequencing performed by Genome Technologies at the Ontario Institute for Cancer Research and Applied Biosystems (Burlington, ON) using the AB SOLiD platform using mate-pair and fragment libraries. For mate-pair libraries, 100 μg of gDNA was sheared to 1-2kb fragments using the GeneMachine HydroShear standard shearing assembly and 1.5kb fragments isolated using 1% agarose gel size selection. Mate-pair libraries were circularized and constructed according to standard SOLiD Long Mate-Pair library protocols (Applied Biosystems, Burlington, ON). Following library quantitation via TaqMan qPCR (Applied Biosystems, Burlington, ON), libraries underwent emulsion PCR and bead enrichment according to standard SOLiD protocols (Applied Biosystems, Burlington, ON). Enriched libraries were then sequenced 2×50bp using SOLiD 3 sequencing chemistry (Applied Biosystems, Burlington, ON). A similar procedure was used to produce fragment libraries, with the following changes: 1 μg of gDNA was sheared to 70-90bp fragments using the Covaris S220 (Covaris Inc., Woburn, MA) and 150bp fragment libraries were constructed according to standard SOLiD Fragment library protocols (Applied Biosystems, Burlington, ON). Libraries were quantified and enriched as described above and sequenced 1×50bp using SOLiD 3 sequencing chemistry (Applied Biosystems, Burlington, ON).

### Sequence alignment and variant calling

Raw reads were split into manageable chunks (n = 10,000,000 reads) and aligned to the rat reference genome (rn6) using BFAST (v0.7.0a) with default parameters. Resulting SAM format files were converted to BAM format and coordinate sorted, followed by mark duplicates, merging of partial files and indexing using Picard (v1.92). Indel realignment/recalibration were performed using GATK (v3.7.0), followed by variant detection using GATK’s HaplotypeCaller, with known variants identified using dbSNP (build 149, downloaded from UCSC on 2018-11-19). Resulting variants were filtered such that any variants with a depth <6 reads and SNPs common to previously sequenced rat strains [28] were excluded [GATKs LiftoverVariants; using rn4 to rn6 chain file from UCSC, and vcftools (v0.1.15)]. Variant annotation was performed using SnpEff (v4.3t) with the rn6 database (Rnor_6.0.86). As these are highly controlled strains, only homozygous variants were carried forward for analysis. This reduced the number of variants by 95% to 176578 for H/W and by 92% to 161620 for L-E.

### Transcription factor binding site analyses

The modified rat genomes (H/W and L-E) were used to identify differing transcription factor binding sites (TFBS) between strains. Conservation scores, REFLINK and REFFLAT tables for rn6 were downloaded from the UCSC genome browser on August 09, 2018 [60]. For each rat strain examined (H/W and L-E), the unique variants identified above were inserted into the reference genome (rn6) FASTA file. For each rat strain, as well as for mouse (mm9), the genome was searched for the following motifs: AHRE-I (core) GCGTG [61], AHRE-I (extended) T/NGCGTG [62], AHRE-I (full) [T|G]NGCGTG[A|C][G|C]A [43] and AHRE-II CATG(N6)C[T|A]TG [63] and ARE TGAC(N3)GC [64]. Exposed motifs were annotated to specific genes if they occurred within a promoter region (±3 kbp of the transcription start site) and a PhyloHMM conservation score from 0 (weak conservation) to 1 (strong conservation) was calculated (**S2 Table**).

## Acknowledgements

The authors thank Hanbert Chen, Alexander Wu, Ashley Smith, Janne Korkalainen, Arja Moilanen, and Virpi Tiihonen for excellent technical assistance and support.

## Supporting information

**S1 Table. Study design.** After outlier removal, a total of 115 animals were used in this study. The number of animals in each experimental group (strain/AHR isoform/TCDD dose) is listed, along with information regarding animals used in comparison analyses from existing studies. Animals were all adult males with all control animals dosed corn oil vehicle.

**S1 Fig. Identification of outliers.** Intensity values for each of 12 “AHR-core” genes following RMA normalization for each array. Red dots indicate outliers as determined by Dixon’s Q test (p < 0.05 for 2 or more genes). A) Experiment #1; three arrays (all from the rWT cohort) were identified as outliers. B) Experiment #2; two arrays (one DBA/2 control and one rWT low dose) were identified as outliers.

**S2 Fig. Microarray QA/QC (Experiment #1).** Quality control analyses of microarrays (post-outlier removal) for samples obtained from the dose response of transgenic mice with C57BL/6 controls. Distributional homogeneity of arrays was assessed both (A) pre-and (B) post-normalization. (C) RNA degradation plots. (D) Inter-array correlation using Pearson’s similarity metric. Similar QA/QC analyses were performed for (E-H) rWT and (I-L) DBA/2 mice.

**S3 Fig. Microarray QA/QC (Experiment #2).** Quality control analyses of microarrays (post-outlier removal) for samples obtained from the dose response of transgenic mice with C57BL/6 controls. Distributional homogeneity of arrays was assessed both (A) pre-and (B) post-normalization. (C) RNA degradation plots. (D) Inter-array correlation using Pearson’s similarity metric. Similar QA/QC analyses were performed for (E-H) rWT, (I-L) DEL and (M-P) INS mice.

**S4 Fig. Overview of transcriptomic profiles.** A) ComBat corrected RMA normalized intensity levels of probesets that demonstrated a variance > 1 across all cohorts (EXP1 and EXP2 combined). B) Samples tended to cluster by treatment (TCDD or corn oil control; ARI = 0.96), rather than genotype or exposure time. Linear modeling was performed on each group (EXP/AHR isoform) separately using RMA normalized values. For EXP2, legend indicates dose of TCDD (125, 250, 500 and 1000 μg/kg) relative to corn oil treated animals: C) C57BL/6 mice, D) DEL, E) INS and F) rWT cohorts. For EXP1, legend indicates dose of TCDD (5, and 500 μg/kg) relative to corn oil treated animals: G) C57BL/6 mice, H) rWT and I) DBA/2 cohorts.

**S5 Fig. Overlap response among cohorts.** Overlap of genes showing significantly altered mRNA abundance (|log_2_ fold change| > 1, p_adj_ < 0.05) among A) C57BL/6, rWT and DBA/2 mice, treated with 5 μg/kg for 19 hours or AHR-ratonized mouse cohorts following exposure to B) 125 C) 250 or D) 1000 μg/kg TCDD for 4 days. Using the same significance criteria, overlap among orthologous gene sets for E) TCDD-sensitive cohorts (500 μg/kg [mouse] or 100 μg/kg [rat], 19 hour exposure), and F) AHR-ratonized mice and corresponding rat strains (500 μg/kg [mouse] or 100 μg/kg [rat], 4 day exposure) are shown.

**S2 Table. Transcription factor binding site analysis.** Orthologous genes were annotated with the presence, absence and conservation score of various AHRE motifs, for both species (mm9 and rn6 genome builds) and rat strains (H/W and L-E, based on the rn6 genome build). In total, 10,706 genes had at least one difference in TFBS between species, and 1,392 genes were shown to differ between H/W and L-E rats in the number of identified TFBS within their promotor regions for at least one of the examined motifs.

**S3 Table. High impact variants unique to Han/Wistar rat.** Whole-genome sequencing of H/W and L-E rats was performed and the resulting variants were annotated as described in methods. 642 high impact, homozygous variants (35 SNVs, 608 indels) were uniquely identified in the TCDD-resistant H/W rat.

**S4 Table. Pathway analysis (mouse-rat comparison).** Orthologous genes between mouse and rat were identified. Significantly altered genes from each indicated cohort (500 μg/kg TCDD for EXP1 and EXP2, 100 μg/kg TCDD for rat cohorts) were submitted for pathway analysis using GoMiner software. Gene Ontologies present in both species were examined. Enrichment and q-value (FDR-adjusted p-value) for each cohort are shown.

**S6 Fig. Transcriptomic profiles for unique Han/Wistar variants.** Whole-genome sequencing of H/W and L-E rats was performed and the resulting variants were annotated as described in methods. 642 high impact, homozygous variants were identified unique to the TCDD-resistant H/W rat. Transcriptomic data was available for 209 of these genes. RNA abundance results for EXP1 (C57BL/6, rWT-mouse and DBA/2: 19 hour exposure to 5 or 500 μg/kg TCDD), EXP2 (C57BL/6, DEL, INS and rWT mice: 4 day exposure to 125, 250, 500 or 1000 μg/kg TCDD) and various rat strains (L-E and H/W: 1, 4, and 10 day exposure to 100 μg/kg TCDD; Ln-A and Ln-C rats: 19 hour exposure to 100 μg/kg TCDD) are shown. Dot size represents the magnitude of change relative to control animals while colour indicates direction of change (orange = increased, blue = decreased); background shading displays significance level (FDR-adjusted p-value). Top covariates show sample information, including AHR and hypothesized gene “B” genotype (NOTE: H/W rats generally express a mixture of INS and DEL AHR variants [predominantly the INS variant]); covariates along the left indicate the predicted consequence of the mutation(s) detected in each gene. Genes with missing data (not included on the microarray used for a given experiment) are indicated by an X.

**S5 Table. Hypergeometric testing.** Hypergeometric testing was performed to determine whether genes demonstrating a novel or lost AHRE-1 (full) motif within their promoter region were more likely to show altered mRNA abundance following TCDD exposure than chance alone (H/W and L-E rats exposed to 100ug/kg TCDD for 4 days); no significant enrichments were found.

**S6 Table. Results of linear modeling (mouse-rat comparison).** Linear modeling was performed to compare treated with control animals for each cohort. Orthologous genes between mouse and rat were identified and results from the two species combined. The magnitude of difference between treatment groups (M) and FDR-adjusted p-value (Q) are shown for each cohort.

